# A Timescale for the Radiation of Photosynthetic Eukaryotes

**DOI:** 10.1101/2020.04.18.047969

**Authors:** Eliane Evanovich, Patricia Jeanne de Souza Mendonça-Mattos, João Farias Guerreiro

## Abstract

Oxygenic photosynthesis is considered the most important evolutionary innovation in the history of Earth. It depends on two photosystems, responsible for the photolysis of water and the reduction of carbon dioxide. Oxygen and carbohydrates are released at the end of the reaction. Extraordinary, the oxygen released created the stratospheric ozone layer, and transformed the ocean chemistry, whereas the carbohydrates are the primary source of energy for complex cells. Several lines of evidence indicate the photosynthesis arose in the ancestors of cyanobacteria. It was spread over some eukaryotes by the acquisition of a free-living cyanobacterium, which evolved into photosynthetic plastid, the chloroplast. The timing of the chloroplast emergence is still controversy. Estimated ages range from 600 to 2100 million years ago (Mya) in accordance to previous studies. The aim of this study is to clarify several aspects of the origin and diversification of photosynthetic eukaryotes. For this purpose, we utilized a data set based on 27 protein-coding genes from genomes of cyanobacteria and photosynthetic eukaryotes, more genes than other papers that also utilized plastid genes, and performed the Bayesian analysis method to estimate the divergence times of the photosynthetic eukaryotes. Results showed photosynthetic eukaryotes emerged Late Mesoproterozoic about 1342 Mya. The Early Proterozoic oceans did not have adequate conditions for eukaryotes, because chemical elements such as zinc and molybdenum were at reduced concentrations, and they are essential to the formation of eukaryotic proteins.

## 1. Background

Oxygenic photosynthesis is probably the most important metabolic innovation of the Earth history. It produces, at the end of the process, water, oxygen and carbohydrates. Remarkable, free oxygen created the stratospheric ozone layer, and changed the ocean chemistry, while the carbohydrates are the main source of energy for complex cells [1, 2, 3, 4]. Most living beings, save for some exceptions, depend directly or indirectly of the oxygenated photosynthesis.

In the Archean Eon, the terrestrial atmosphere and the oceans were anoxic. This began to change about 2500 million years ago (Mya) after the Great Oxygenation Event (GOE). Later, four more oxygenation events happened in the primitive Earth. In the Proterozoic from 1850 to 850 Mya, the oceans became slightly less anoxic and sulfidic [5, 6, 7, 8]. These environmental changes appear to have triggered an increase in Proterozoic biodiversity, even from the complex eukaryotes carrying specialized structures [6, 8].

Several studies suggest that the oxygenic photosynthesis arose in the ancestors of Cyanobacteria (blue-green algae), and it was spread over some eukaryotes, through the acquisition of a free-living cyanobacterium, which evolved into photosynthetic plastid, the chloroplast [1, 4, 9]. This event, known as primary endosymbiosis arose in Archaeplastida, a monophyletic group formed by Rhodophyta, Viridiplantae and Glaucophyta [10, 11]. In contrast to Archaeplastida, CASH organisms (Cryptophyceae, Alveolata Stramenopiles, and Haptophyta) are paraphyletic, however, share a red plastid, which was obtained by secondary endosymbiosis with a red alga. CASH plastids (or complex red plastids) have the same protein transport apparatus, and present four membranes [13, 14, 15, 16, 17, 18, 19]. Conforming to the hypothesis of a unique origin of the red plastids, the loss of plastids in groups as some alveolates (apicomplexans and dinoflagellates), *Goniomonas* (Cryptophyceae), and Stramenopiles (Bicosoecida, Labyrinthulomycetes, Oomycetes, and Opalinidae) would be a derived condition [20, 21]. In accordance with the taxonomic review for Eukarya [22], Stramenopiles, Alveolata and Rhizaria form the SAR clade, whereas Cryptophyceae and Haptophyta do not have established phylogenetic positions within the Diaphoretickes (Domain which comprises Cryptophyceae, Haptophyta, Archaeplastida, SAR, and Euglenozoa). Dinoflagellate plastid *Durinskia* and *Kryptoperidinium* likely were retrieved from diatoms, while the Chlorarachniophyceae plastid derived from a Trebouxiophyceae alga by tertiary endosymbiosis [23, 24].

Chloroplast DNA (cpDNA) has about one-twentieth the size of cyanobacterial genomes [25]. It is circular or linear and includes genes involved in transcriptional machinery (ribosomal RNA, transfer RNA, and RNA polymerase genes), photosynthesis, and biosynthesis of molecules as vitamins, amino acids, pigments, and fatty acids [25, 26]. The origin of the chloroplast is still a controversial issue. Previous studies assign an older aged between 1423 to 2100 Mya [27, 28, 29, 30, 31], while others have estimated a younger aged between 600 to 1250 Mya [32, 33, 34, 35]. Results found by Betts et al. [35] for the divergence time among primary endosymbiosis showed larger limits, between 1118 to 1774 Mya. Estimates of divergence times cited above were based on molecular data, except for the one proposed by Cavalier-Smith [33] which was based on fossil records and their geochemical impacts. In this paper, we performed relaxed Bayesian estimates based on 27 plastid genes to infer the possible date of origin and diversification of photosynthetic eukaryotes. Compared to dataset utilized by Yoon [28], we used more genes (21 more genes) and algae groups (Chlorarachniophyceae and Dinoflagellata), and found an age of 1342 Mya for the origin of photosynthetic eukaryotes.

## 2. Methods

### 2.1. Samples and Alignment

Genome sequences utilized in this paper were available from the National Center for Biotechnology Information (NCBI) (https://www.ncbi.nlm.nih.gov). Description and source of the samples are on additional file: Table S1. MUSCLE software [36] was utilized to perform sequence alignment of 27 protein-coding genes (additional file: Table S2). The concatenated sequence had 23240 nucleotides and was later submitted to Gblocks 0.91b [37], which selected the most conserved matrix data. Gblocks generated an alignment containing 11606 ungapped nucleotides, 49% of the original sequence.

### 2.2. Molecular Dating Analyses

GTR+I+G was the most appropriate model and was selected by Jmodeltest 2.1.10 [38]. RaxML 8 software [39] was applied to generate phylogenies. The best tree was utilized as an input tree on BEAST. Phylogenies obtained are in the additional file: Dataset S1. BEAST v2.6.0 [40] was performed to measure the divergence dates. The analysis was based on the following models: GTR+I+G, gamma-distributed heterogeneity of substitution rates among sites, uncorrelated log-normal relaxed clock (UCLN), and Yule. The computational performance was improved by BEAGLE library [41]. We run five independent Metropolis-coupled Markov chain Monte Carlo (MCMCMC) in BEAST. Each had a chain of 100 million generations and burn-in of 10 million. Tracer 1.7 [42] was utilized to monitor the analyzes and indicate when there was the convergence. At the end of the analysis, 100 trees were generated. TreeAnnotator [43] was applied to measure the consensus tree that was viewed with FigTree v1.4.4 [44]. The fossil calibration points are shown in Table 1.

**Table 1.** Description of the fossils utilized as calibration points in this study.

| **Taxon** | **Fossil** | **Calibration point (Mya)** | **Type constraints** | **Reference** |
| --- | --- | --- | --- | --- |
| Cyanobacteria | Possible microfossils | 3500 | Maximum | 45 |
| Cyanobacteria | Biomarker evidence | 1640 | Minimum | 46 |
| Eukarya | *Grypania spiralis* | 2100 | Maximum | 47 |
| Eukarya | Possible protist microfossils | 1500 | Minimum | 48 |
| Rhodophyta: Bangiophyceae | *Bangiomorpha pubescens* | 1047 | Minimum | 34, 49 |
| Viridiplantae: Chlorophyta | *Proterocladus antiquus* | 1000 | Maximum | 50 |
| Stramenopiles: Xanthophyceae | *Palaeovaucheria* | 1000 | Maximum | 51 |
| Stramenopiles: Xanthophyceae | *Jacutianema* | 750 | Minimum | 52 |
| Rhodophyta: Florideophyceae | Oldest known Florideophyceae fossil | 550 | Minimum | 48 |
| Stramenopiles: Phaeophyceae | *Punctariopsis latifolia* | 490 | Minimum | 53 |
| Haptophyceae | First appearance of calcareous nannofossil | 225 | Minimum | 54, 55 |
| Viridiplantae: Zygnemophyceae | *Tetraporina horologia* | 345 | Maximum | 56 |
| Viridiplantae: Zygnemophyceae | *Paleoclosterium spiralis* | 93 | Minimum | 57,58 |
| Stramenopiles: Bacillariophyta | Oldest known diatom | 185 | Minimum | 59 |
| Stramenopiles: Bacillariophyta | Pennate diatoms | 50 | Minimum | 60 |

## 3. Results and Discussion

### 3.1. Phylogenetic Relationships

Topology found in this paper was similar to that obtained by Janouškovec et al. [18], which also utilized chloroplast genes. Phylogenetic trees based on chloroplast genes have often failed to recover the Archaeplastida monophyly because they reflect more recent phylogenetic relationships containing algae arising from secondary and tertiary endosymbiosis. Figure 1 shows the time scale for the origin and diversification of the photosynthetic eukaryotes. Phylogenetic tree with the minimum and maximum limits is shown in the additional file: Figure S1.

**Figure 1:**
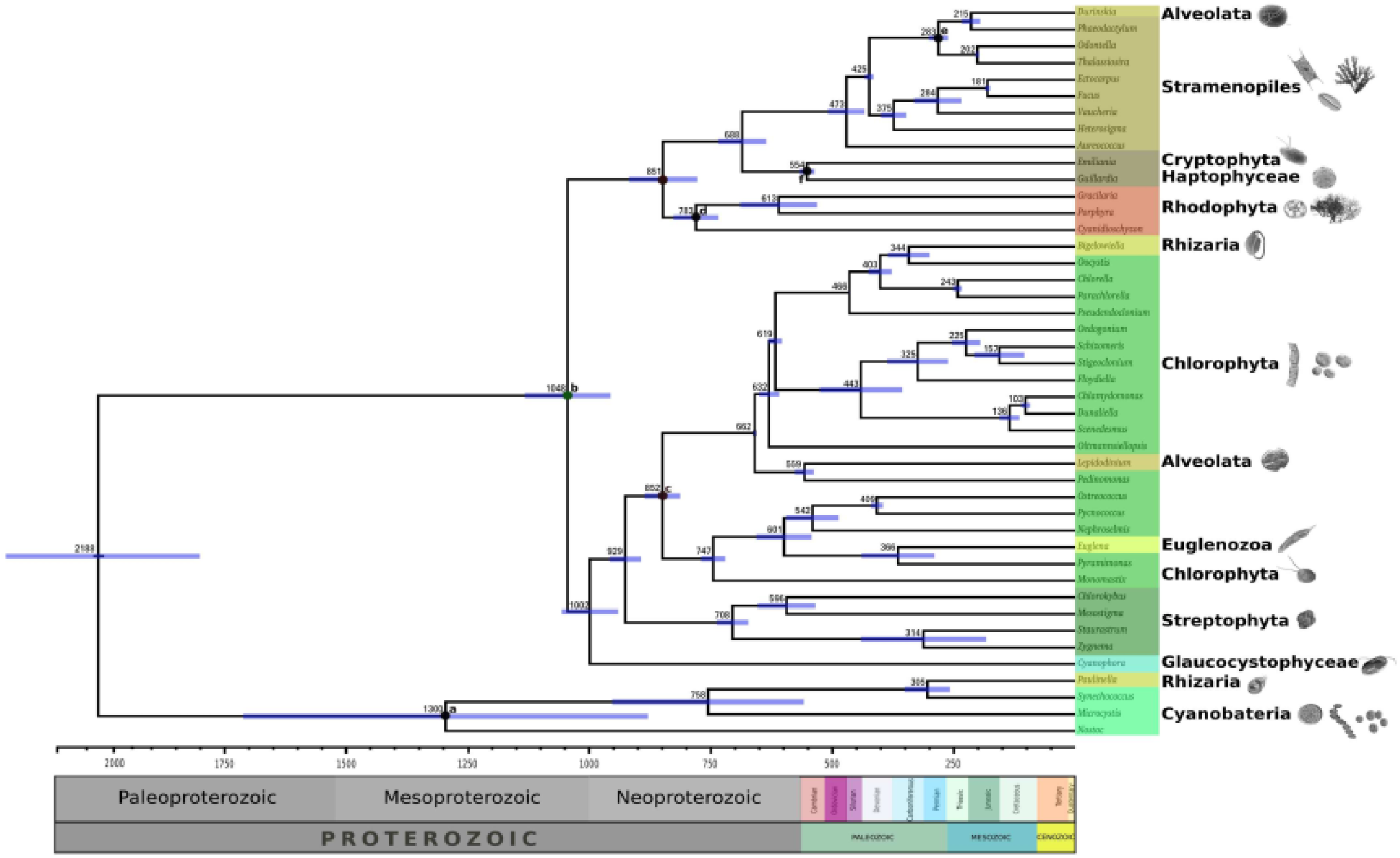
A time scale for the origin and diversification of the photosynthetic eukaryotes obtained by BEAST 2 based on 27 chloroplast-encoded genes. The estimated mean divergence times are shown at the nodes of the tree, and the blue bars at nodes represent the 95% confidence interval. Calibration nodes are indicated by the circles. Dark circles correspond to nodes with two calibration points, and light circles are nodes with one calibration point. Above of the tree is presenting the geological time.

### 3.2 Primary Plastid Endosymbiosis

Origin of the chloroplast was about 1342 Mya (95% highest posterior density or HPD: 1155-1848 Mya) according to our results, and it is closer to limits estimated by Betts et al. [35], and Yoon et al. [28]. Gibson et al. [34] rejected *Bangiomorpha* as representative of Bangiales-Rhodophyta. Our results do not reject the phylogenetic position of these fossils and other Proterozoic fossils such as *Proterocladus antiquus, Palaeovaucheria*, and *Jacutianema*.

It is noteworthy that the low oxygen levels in the Paleoproterozoic (2500 to 1600 Mya ago) were incompatible with the bioavailability of essential micronutrients for the enzymatic activity of eukaryotic cell [5, 6]. In addition to having low levels of micronutrients, the Paleoproterozoic oceans also had high sulfate concentrations [61, 62, 63]. The biogeochemical stasis previously experienced by the planet was finally broken in Mesoproterozoic [64], and abrupt innovations appeared on Earth for the first time during the Neoproterozoic (1000 to 541 Mya ago) as ice sheets, alteration in carbon and oxygen levels, and increase in primary productivity [64]. The emergence and diversification of eukaryotic algae may probably have had to play a decisive role in these transformations.

### 3.3. Glaucophyceae

In Archaeplastida, Glaucophyceae was the first lineage to emerge about 1322 Mya (95% HPD: 1103-1824 Mya), shortly after the emergence of photosynthetic eukaryotes. These algae retain primitive characters, similar to an ancestral cyanobacterium, such as a vestigial peptidoglycan wall, non-stacked thylakoid, and carboxysome-like bodies, therefore it is considered a living fossil [65]. It has little diversity, restricted to four genera (Cyanophora, Cyanoptyche, Glaucocystis, and Gloeochaete), which contrasts with the vast diversity found in Rhodophyta and Viridiplantae.

### 3.4. Viridiplantae

The divergence between Streptophyta and Chlorophyta was estimated at 1108 to 1401 Mya (mean 1244 Mya), whereas the emergence of Chlorophyta was between 984 to 1203 Mya (mean 1093 Mya). This knowledge is consistent with a possible Chlorophyta fossil called *Proterocladus antiquus* dated 1000 Mya [50]. Radiation of Viridiplantae may have influenced the fluctuations in the carbon cycle and consequently would have helped to trigger the initiation of Snowball Earth [64].

### 3.5. Rhodophyta

The age estimated for the Rhodophyta radiation was 1127 Mya (95% HPD: 1061-1242 Mya), which is consistent with the age of 1047 Myr[34]. In according to Cavalier-Smith [64], *Bangiomorpha pubescens* would be a bacterial complex, whereas Berney and Pawlowski [33] and Gibson et al. [34] suggest that *Bangiomorpha* might be a stem photosynthetic eukaryotic extinct. The large size, sexual reproduction, multicellularity, and radial cell division are the traits that supported *Bangiomorpha* as a member of the Rhodophyceae [49].

### 3.6. Secondary Plastid Endosymbiosis

Secondary endosymbiosis at least three types of plastids, found in Euglenozoa, Chlorarachniophyceae, and CASH organisms [66, 67]. Results show the engulfment of Pyramimonadales algae by Euglenozoa ancestor occurred about 366 Mya in the Upper Devonian. Oldest known fossil of Euglenozoa dates from Lower Cretaceous [68, 69]. Chlorarachniophyte plastids were originated from a Trebouxiophyceae alga [24] during the Carboniferous (344 Mya). Our estimates indicate the red plastid emerged about 1156 Mya (95% HPD: 1070-1313) during the Neoproterozoic. Stramenopiles arose between 1044-1182 Mya (mean 1017 Mya). The date is within the range estimated by Berney and Pawlowski [33]. And Coccolithophores (Haptophyta) would have appeared about 538 to 570 Mya (mean 554 Mya). These algae have a fundamental role in the production and precipitation of calcium carbonate (CaCO3) and besides influence the light scattering and cloud cover from the Jurassic [70].

### 3.7. Tertiary Plastid Endosymbiosis

Some Peridiniales plastids were obtained from red algae, however, none of them is represented in this paper. Our analysis was restricted to *Durinskia* and *Kryptoperidinium*. Their plastids were acquired from the engulfment of a diatom between 64.5 to 194 234 Mya (mean 109 Mya). The most reliable fossil record of these algae appear since Triassic, had an increase in Cenozoic Era, and is currently in decline [71].

## 4. Conclusion

The biogeochemical stasis of the Earth lasted about 3 billion years, and it was broken during the Late Mesoproterozoic. Earlier the oceans were inhospitable, in other words, highly sulfidic and poorly oxygenated. Sulfides are toxic to eukaryotic algae, and the low levels of oxygen and micronutrients inhibit the formation of proteins in these organisms. Our results indicate that photosynthetic eukaryotes arise around 1048 Mya, at the end of Mesoproterozoic, and diversified rapidly. The radiation of photosynthetic eukaryotes helped progressively change the chemical composition of the oceans, increased the primary productivity and favoured the emergence of new life forms. Early eukaryotes may have had a wide variety of organisms, such as *Bangiomorpha* and *Paleovaucheria*. Many of them have disappeared or have a yet unknown phylogenetic affinity, similar to what happened too many representatives of the Ediacaran biota.

## Abbreviattions

CaCO_3_: calcium carbonate
cpDNA: Chloroplast DNA
GOE: Great oxygenation event
HPD: 95% Highest posterior density
LECA: last eukaryotic common ancestor
MCMCMC: Metropolis-coupled Markov chain Monte Carlo
MRCA: most recent common ancestor
Mya: million years ago
NCBI: National Center for Biotechnology Information
PRONE: plant-specific Rop nucleotide exchanger
Re-Os: Rhenium-Osmium
rRNA: ribosomal RNA
tRNA: transfer RNA

## Data Availability

Supporting data are enclosed as additional files.

## Additional Points

This manuscript is part of the Ph.D. thesis of Eliane Evanovich from Programa de Pós-graduação em Genética e Biologia Molecular, Universidade Federal do Pará. The authors wish to express their thanks for the financial support of Pró-Reitoria de Pesquisa e Pós-Graduação (PROPESP).

## Conflicts of Interest

The authors declare that they have no competing interests.

### Authors’ contributions

The authors read and approved the final manuscript.

